# Alpha Phase Dynamics Predict Age-Related Visual Working Memory Decline

**DOI:** 10.1101/050450

**Authors:** Tam T. Tran, Nicole C. Hoffner, Sara C. LaHue, Lisa Tseng, Bradley Voytek

## Abstract

Alpha oscillations are modulated in response to visual temporal and spatial cues, However, the neural response to alerting cues is less explored, as is how this response is affected by healthy aging. Using scalp EEG, we examined how visual cortical alpha activity relates to working memory performance. Younger (20-30 years) and older (60-70 years) participants were presented with a visual alerting cue uninformative of the position or size of a lateralized working memory array. Older adults showed longer response times overall, and reduced accuracy when memory load was high. Older adults had less consistent cue-evoked phase resetting than younger adults, which predicted worse performance. Alpha phase prior to memory array presentation predicted response time, but the relationship between phase and response time was weaker in older adults. These results suggest that changes in alpha phase dynamics, especially prior to presentation of task-relevant stimuli, potentially contribute to age-related cognitive decline.

---

In order to achieve high behavioral performance, limited attentional resources must be efficiently directed towards task-relevant information. Such information could include the timing or spatial position of upcoming visual stimuli. Knowledge of when^1^ or where^2^ a target will appear enhances detection and shortens response times. Likewise, presentation of neutral warning cues improves response times by heightening alertness or preparedness for upcoming stimuli. The effects of informative temporal and spatial cues are strongly related to the dynamics of 7-14-Hz alpha oscillations, as observed in anticipatory changes in alpha amplitude^3–6^ and phase^7^. How alpha dynamics are modulated in response to noninformative alerting cues is less understood.

Neurologically healthy aging is associated with declines in attention and working memory. Behaviorally, the benefits of spatial cuing are relatively resistant to healthy aging^8,9^, but older adults derive less benefit from the presence of temporal^5^ and alerting cues^10,11^. Physiologically, older adults show reduced alpha modulation in response to temporal^5^ and spatial cues^12^, though a recent study found no age-related differences in neural response to alerting cues^13^. However, because alpha activity was not examined in that study, it is unclear whether older adults’ reduced use of alerting cues can be predicted by concomitant changes in alpha oscillatory dynamics.

To investigate alpha response to alerting cues, and how this response is affected by healthy aging, we recorded EEG from younger and older adults performing a unilateral visual working memory task. Each trial of the task included an alerting cue signaling the upcoming presentation of a lateralized memory array. This cue allowed us to probe participants’ preparedness for upcoming stimuli independent of motor preparation. We hypothesized that age-related changes in neural activity would manifest themselves in the alpha amplitude and phase response to presentations of the alerting cue. We also hypothesized that the extent to which neural response to the alerting cue was altered would also predict declines in working memory performance.

## Results Behavior

### Response Time

We compared younger and older adults’ response times (RTs) on a lateralized visual working memory task (Fig. 1a, see Methods). RTs showed main effects of age (Fig. 1b, *F*_1,29_ = 13.32, *p* = 0.0010, generalized η^2^ = 0.31) and memory load (*F*_2,58_ = 67.20, Greenhouse-Geisser (GG) ε = 0.88, *p*_GG_ < 10^−13^, η^2^ = 0.089) and an interaction between age and memory load (*F*_2,58_ = 3.75, ε = 0.88, *p*_GG_ = 0.029, η^2^ = 0.0054). Between groups, younger adults had faster RTs than older adults in each load condition. This included load-one (541 ms vs. 643 ms, mean difference 95% confidence interval [-166 ms, −39 ms], *t*_28.91_ = −3.29, *p* = 0.0027, Cohen’s *d* = −1.17), load-two (565 ms vs. 670 ms, [-166 ms, −44 ms], *t*_29_ = −3.51, *p* = 0.0015, Cohen’s *d* = −1.24), and load-three conditions (591 ms vs. 721 ms, [-195 ms, −65 ms], *t*_29_ = −4.09, *p* < 10^−3^, Cohen’s *d* = −1.45).

**Figure 1.**
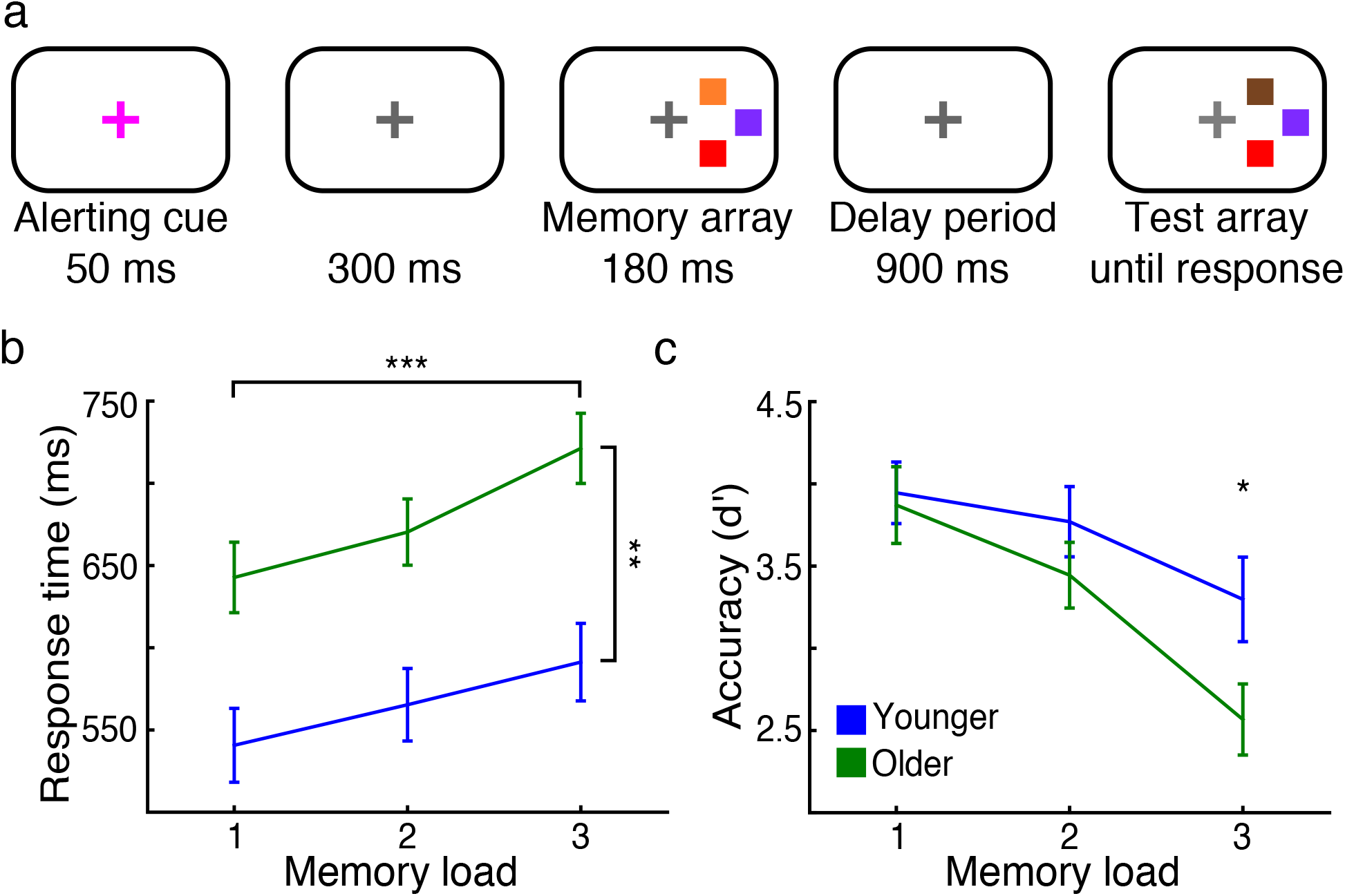
Paradigm, behavioral performance, and event-related activity. (*a*) Diagram of the task design, in this example showing a non-matching test array. (*b*) Response times increased with increasing memory load, with younger adults (blue) faster than older adults (green, \*\**p* < 0.01, \*\*\**p* < 0.001; error bars, SEM). (*c*) Accuracy decreased with increasing memory load, with younger adults more accurate than older adults during load-three trials (\**p* < 0.05; age by memory load interaction: *p* < 0.01; error bars, SEM).

### Accuracy

As assessed using the sensitivity measure *d’*, accuracy showed an effect of memory load (Fig. 1c, *F*_2,58_ = 51.04, ε = 0.92, *p*_GG_ < 10^−11^, η^2^ = 0.16) and an interaction between age and memory load (*F*_2,58_ = 5.78, ε = 0.83, *p*_GG_ = 0.0065, η^2^ = 0.021). Accuracy was comparable between younger and older adults in load-one (*p* = 0.73, Cohen’s *d* = 0.13) and load-two conditions (*p* = 0.22, Cohen’s *d* = 0.45). However, younger adults outperformed older adults in load-three conditions (3.32 vs. 2.58, [0.042, 1.45], *t*_29.00_ = 2.17, *p* = 0.039, Cohen’s *d* = 0.77). In summary, older adults showed slower RTs overall and reduced working memory accuracy specifically during high-load trials.

## EEG

### Alerting Cue Activity

To investigate neurophysiological measures potentially underlying decreased behavioral performance in older adults, we first examined younger and older adults’ neural response to presentations of the alerting cue. During task performance, younger and older adults exhibited 7-14 Hz oscillatory alpha activity in visual parietal-occipital regions (Fig. 2a). Based on participants’ peak alpha frequency, previously shown to be lower in older adults^14^, we determined individualized alpha bands and compared participants’ normalized alpha analytic amplitude and instantaneous phase activity during the task. To examine the consistency in alpha phase activity across trials, we also computed alpha intertrial coherence (ITC) per participant.

**Figure 2.**
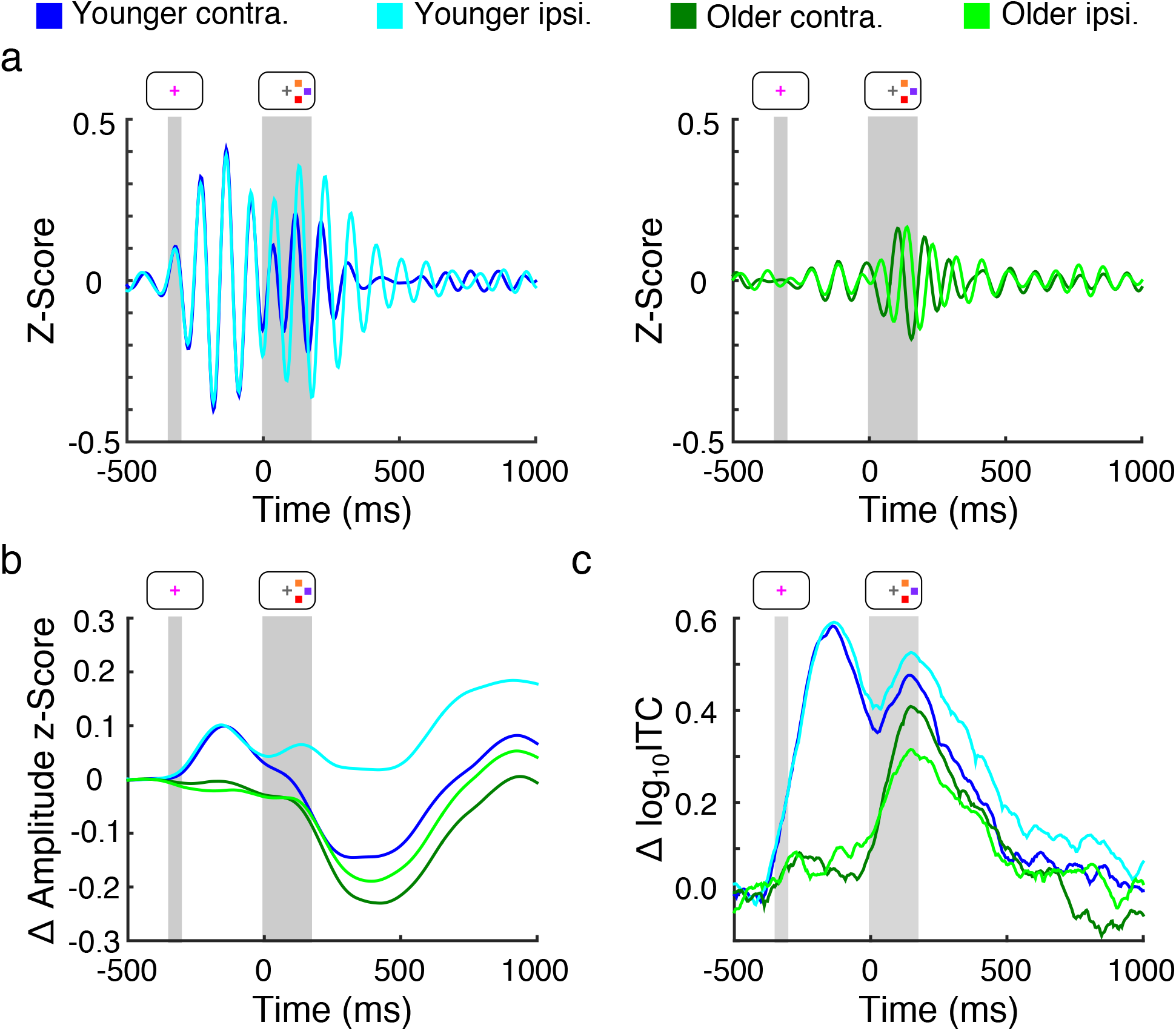
Alpha amplitude and phase activity. a*A*) Grand average visual-area alpha activity contralateral (darker) and ipsilateral (lighter) to the memory array in younger (blue, left panel) and older adults (green, right panel). Gray regions indicate presence and duration of the alerting cue and memory array. Note the hemispheric amplitude differences and strong phase consistency in younger compared to older adults. (*b*) Grand average of changes in visual-area alpha amplitude and (*c*) intertrial coherence relative to baseline, emphasizing the effects observable in (*a*).

Parietal-occipital visual regions showed alpha amplitude and ITC response to presentations of the alerting cue (Fig. 2b, 2c). Alpha amplitude modulation in response to the alerting cue (−350 to 0 ms) showed no effects of age (*F*_1,29_ = 2.82, *p* = 0.10, η^2^ = 0.074), hemisphere (*F*_1,29_ < 1.0), or memory load (*F*_2,58_ < 1.0). This lack of hemisphere and memory load effect is consistent with the alerting cue being uninformative of the lateral position and number of upcoming stimuli.

Compared to baseline (−500 to −350 ms), alpha ITC increased in response to the alerting cue in both younger and older adults. Using a resampling procedure to compare cue-evoked to baseline ITC on a per-subject basis, we determined that all 17 younger adults, as well as 11 out of 14 older adults, showed cue-evoked increases in ITC (*p* < 10^−4^ for all). These increases in ITC suggest the presence of stimulus-evoked alpha phase resets in both younger and older adults. As with alpha amplitude, cue-evoked ITC did not show an effect of hemisphere (*F*_1,29_ < 1.0) or memory load (*F*_2,58_ < 1.0), again consistent with the noninformative nature of the alerting cue. However, younger adults had higher cue-evoked ITC than did older adults (Fig. 3a, 3b, 0.63 vs. 0.23, [0.24, 0.56], *F*_1,29_ = 23.64, *p* < 10^−4^, η^2^ = 0.32).

**Figure 3.**
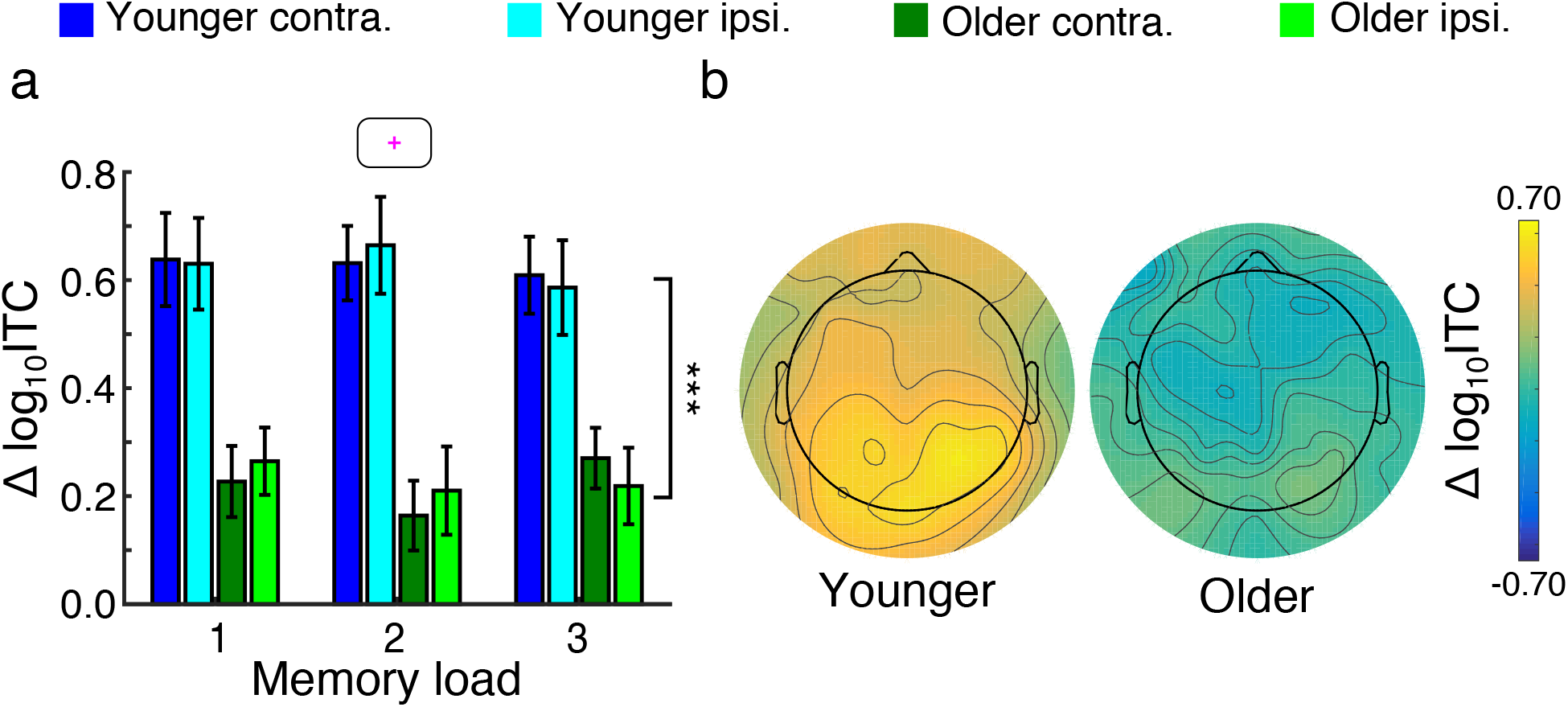
Alerting cue activity. (*a*) Peak alpha intertrial coherence (ITC) in response to the alerting cue. Younger adults (blue) had higher cue-evoked ITC than older adults (green; \*\*\**p* < 0.001; error bars, SEM). (*b*) Topographies of cue-evoked ITC response in younger (left) and older adults (right) during load-three trials.

### Memory Array Activity

Younger adults also showed alpha response to presentation of the memory array. After memory array onset, alpha amplitude diverged between hemispheres in younger and older adults (Fig. 2b). Mean alpha amplitude (0 to 400 ms) showed main effects of memory load (Fig. 4a, 4b, *F*_2,58_ = 4.29, ε = 0.87, *p*_GG_ = 0.024, η^2^ = 0.011) and hemisphere (*F*_1,29_ = 18.15, *p* < 10^−3^, η^2^ = 0.034) and an interaction between age and hemisphere (*F*_1,29_ = 9.10, *p* = 0.0053, η^2^ = 0.017). Post hoc analysis revealed that alpha amplitude decreased from load-one to load-two ([0.0053, 0.056], *t*_30_ = 2.47, *p* = 0.019, Cohen’s *d* = 0.44), but not from load-two to load-three conditions (*p* = 0.37, Cohen’s *d* = 0.17). In addition, alpha lateralization, or the difference in alpha amplitude between hemispheres, was greater in younger than older adults (0.11 vs. 0.019, [0.034, 0.15], *t*_23.21_ = 3.22, *p* = 0.0038, Cohen’s *d* = 1.09).

**Figure 4.**
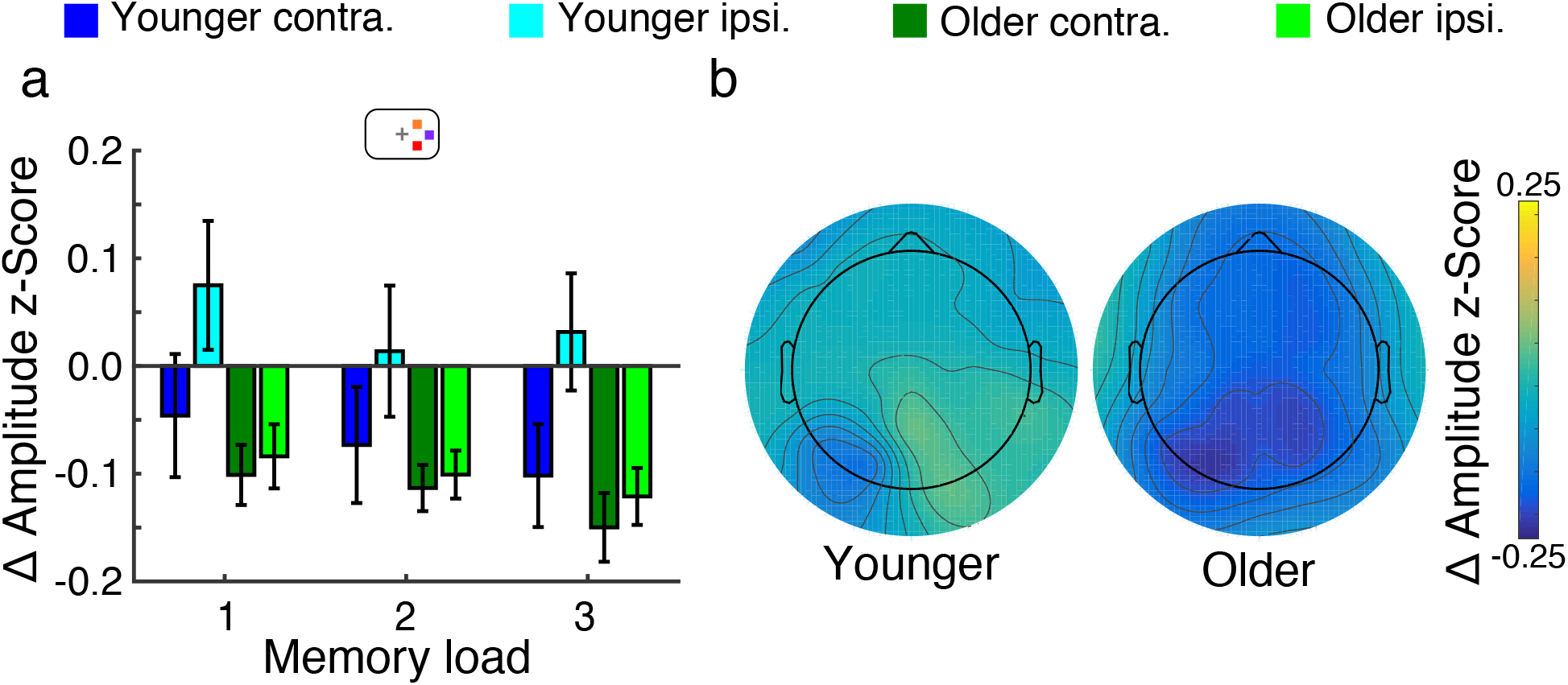
Memory array activity. (*a*) Average change (relative to baseline) in alpha amplitude 0 to 400 ms after memory array presentation. Amplitude decreased from load one to two (*p* < 0.05), and older adults (green) showed decreased alpha lateralization (*p* < 0.01; error bars, SEM). (*b*) Topographies of delay-period alpha amplitude in younger (left) and older adults (right) during load-three trials.

As with alerting cue presentation, memory array presentation also caused alpha phase resets (Fig. 2c). Overall, 15 of 17 younger adults as well as 12 of 14 older adults showed array-evoked ITC (*p* < 10^−4^ for all). Unlike with cue-evoked ITC, array-evoked ITC showed no effects of memory load (*F*_2,58_ < 1.0), age (*F*_1,29_ = 1.60, *p* = 0.22, η^2^ = 0.028), or hemisphere (*F*_1,29_ < 1.0).

### Contralateral Delay Activity

We also investigated participants’ contralateral delay activity (CDA), an event-related potential measure indicative of working memory capacity^15,16^ and top-down attentional processes^17–20^. We observed sustained delay-period (300 to 900 ms) negativity in the hemisphere contralateral to the memory array (Fig. 5a). This negativity or CDA showed a main effect of memory load (Fig. 5b, F_2,58_ = 14.88, ε = 0.96, *p*_GG_ < 10^−5^, η^2^ = 0.080), wherein CDA increased in magnitude from load-one to load-two conditions ([0.34 *μ*V, 0.86 *μ*V], *t*_30_ = 4.66, *p* < 10^−4^, Cohen’s *d* = 0.84). CDA was comparable between load-two and load-three conditions (*p* = 0.47, Cohen’s *d* = 0.13). However, CDA did not differ between younger and older adults (F_1,29_ = 1.05, *p* = 0.31, η^2^ = 0.029), nor did we observe an interaction between age and memory load (F_2,58_ < 1.0).

**Figure 5.**
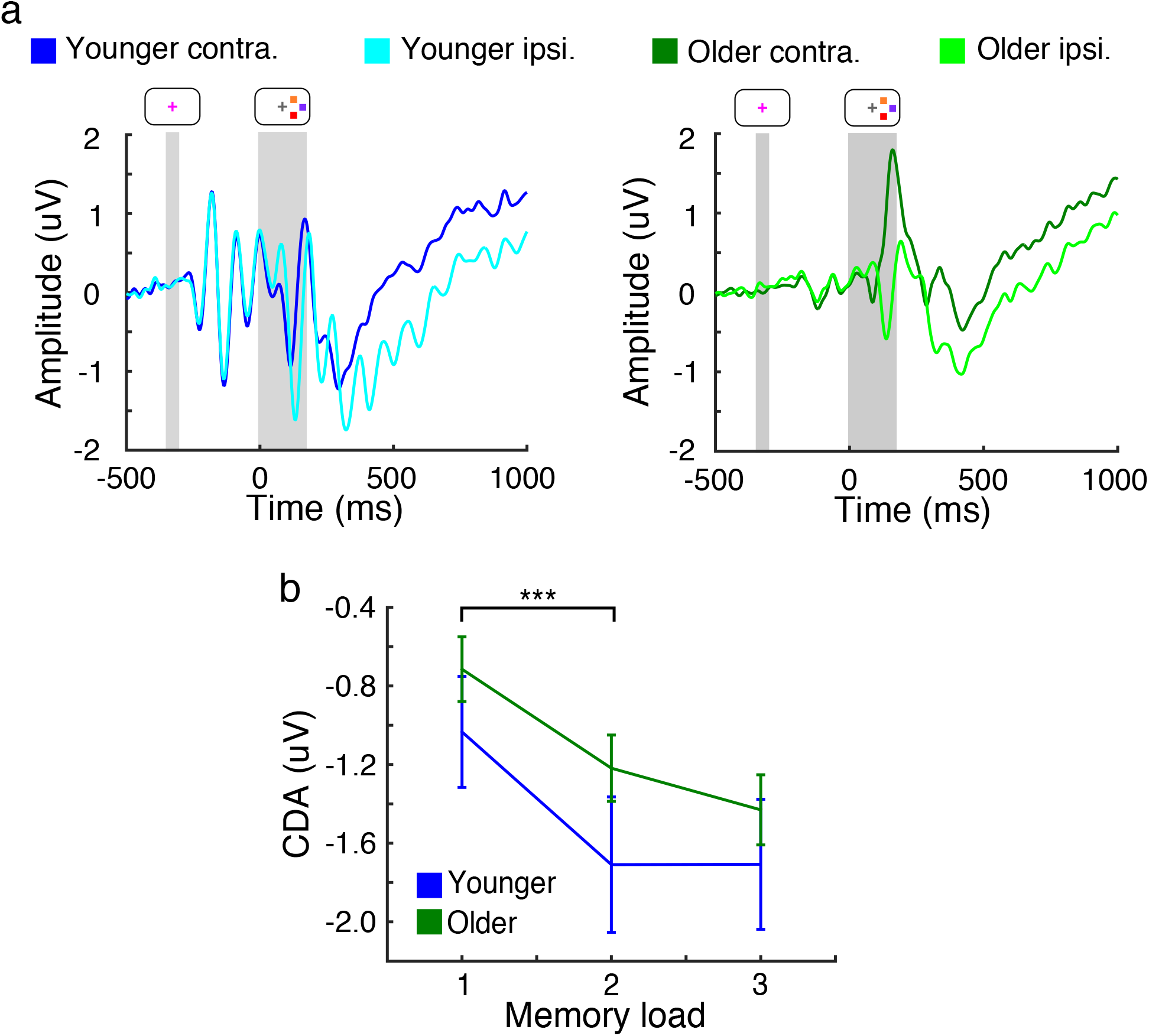
Event-related potential and delay period activity. (*a*) Grand average visual-area activity contralateral (darker) and ipsilateral (lighter) to the memory array in younger (left panel) and older adults (right panel). Gray regions indicate presence and duration of the alerting cue and memory array. Note the sustained negativity in the contralateral hemisphere in both younger and older adults. (*b*) Contralateral delay activity (CDA) increased in magnitude from load-one to load-two conditions, but did not differ between younger and older adults (\*\*\**p* < 0.001; error bars, SEM)

### Alpha Phase Activity Predicts Behavior

Given the age-related changes in neural activity that we observed, we examined how these changes related to behavioral performance. As noted, older adults performed as well as younger adults on the easiest (load-one and load-two) trials, but performed worse for more difficult load-three trials. We examined the neurophysiological basis for this behavioral aging effect using a multiple linear regression analysis. This analysis allowed us to examine the relative contribution of each neurophysiological measure to behavioral accuracy. Specifically, we modeled *d’* as a linear combination of cue-evoked alpha ITC, array-evoked alpha amplitude modulation, and delay-period CDA. Cue-evoked ITC was averaged across visual hemispheres, and the lateralized difference was used for array-evoked amplitude activity. Importantly, these physiological measures were indexed during times *prior to* the actual memory challenge and thus are related to trial-by-trial changes in alertness, encoding, or memory maintenance, rather than memory retrieval or response.

This model explained 17.1% of the variance in accuracy (*p* = 0.045). Examining the relative contribution of each predictor, we found that after accounting for alpha lateralization and CDA, cue-evoked ITC remained predictive of load-three accuracy (*p* = 0.013). Cue-evoked ITC was correlated with load-three accuracy across all participants (Fig. 6a; *N* = 31, Spearman’s *r* = 0.47, *p* = 0.0071) and correlated with load-three accuracy across younger adults alone (*N* = 17, Spearman’s *r* = 0.49, *p* = 0.044). Alpha lateralization and CDA, on the other hand, did not remain predictive of load-three accuracy after accounting for other physiological measures (*p* = 0.28, *p* = 0.94). Thus, cue-evoked ITC prior to the presentation of to-be-remembered stimuli was a strong predictor of behavioral accuracy, even after adjusting for array-related alpha amplitude and delay-period CDA effects.

**Figure 6.**
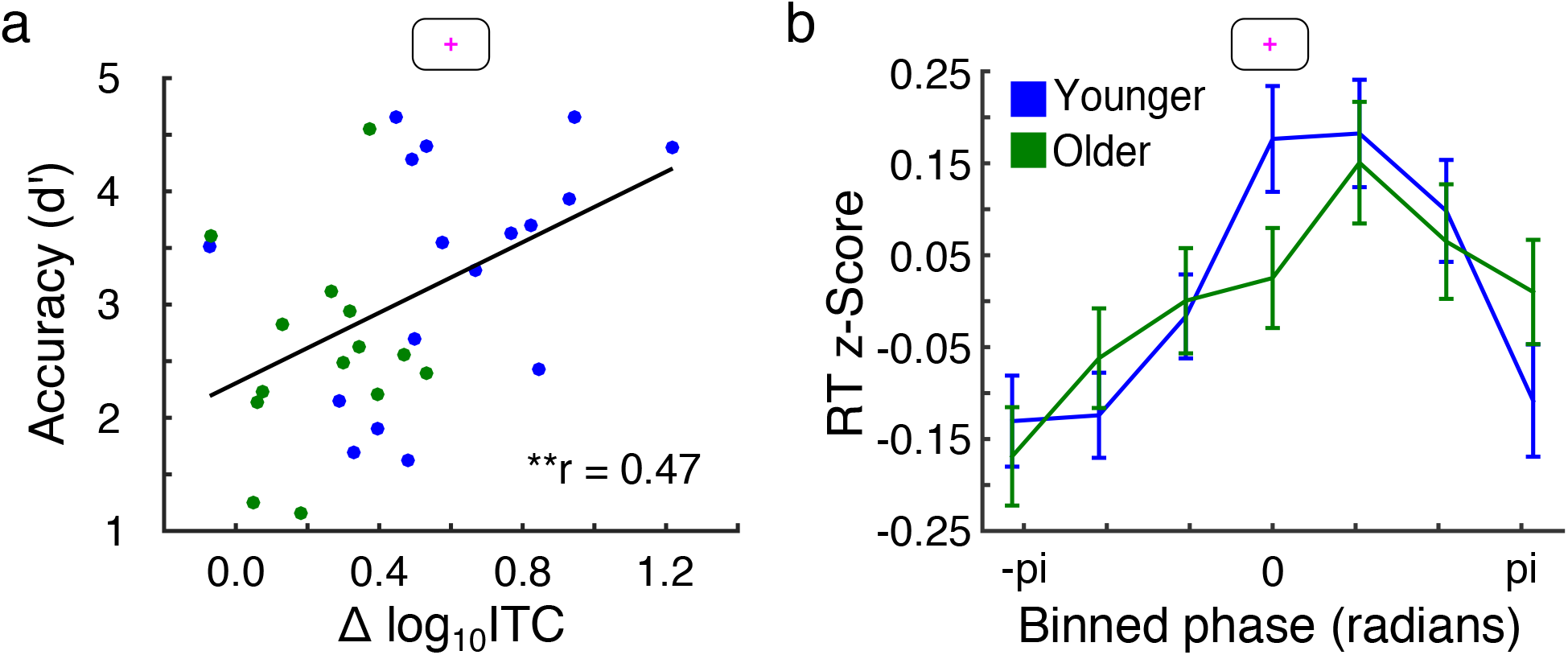
Alpha phase predicts working memory performance. (*A*) Cue-evoked alpha intertrial coherence (ITC) versus accuracy during load-three trials across younger (blue) and older adults (green). Cue-evoked ITC was predictive of load-three accuracy (\*\**p* < 0.01). (*B*) Average response time (RT) binned by alpha phase at peak cue-evoked ITC. Phase of zero and ±pi correspond to the peaks and troughs of alpha, respectively. Trial-by-trial alpha phase predicted RTs (*p* < 10^−3^; error bars, SEM).

To further investigate how cue-evoked alpha ITC is associated with behavioral performance, we examined how alpha phase at peak ITC related to subsequent working memory performance. To do so, we determined the timepoint of each participant’s peak cue-related ITC, and we pooled all participants’ corresponding alpha phases and RTs across trials. During load-three trials in younger adults, alpha phase at peak cue-evoked ITC predicted RTs on a trial-by-trial basis (Fig. 6b, blue; *N* = 2499, *r* = 0.14, *p* < 10^−3^). Alpha phase at peak cue-evoked ITC also predicted RTs in older adults (Fig. 5B, green; *N* = 2090, *r* = 0.079, *p* = 0.0015). Thus, despite older adults’ relatively inconsistent cue-evoked phase response, prestimulus alpha phase was still predictive of load-three RTs. However, the relationship between alpha phase at peak cue-evoked ITC and RT was weaker in older than younger adults (*z* = 1.95, *p* = 0.026).

## Discussion

In this study, we used a combined visual attention and working memory task to investigate how age-related changes in alertness and spatial attention affect later working memory performance. Using scalp EEG, we found that alpha activity showed age-related alterations during the task, including in older adults’ reduced alpha amplitude lateralization during working memory maintenance. In addition, prior to working memory encoding, older participants showed less consistent phase response to a noninformative alerting cue. The consistency of cue-evoked alpha phase reset predicted working memory performance, as did prestimulus alpha phase prior to memory array presentation. Our results provide evidence that alerting cue presentation is accompanied by alpha activity modulation, that neural response to alerting cues is altered during healthy aging, and that the degree of alteration could influence behavioral outcomes.

In this task, compared to younger adults, older adults had slower response times in each memory load condition, but lower accuracy only during load-three trials. These slower response times could indicate a speed-accuracy trade-off strategy in older adults, perhaps accounting for older adults performing with accuracy comparable to that of younger adults in low-load conditions. In addition to their longer response times, older adults were less accurate in high-load trials. Thus, any benefit of slowing was unable to preserve performance in high-load trials, underscoring that age-related reductions in attention and working memory performance are more readily apparent during increasingly difficult tasks.

Previous research has found that contralateral delay activity (CDA) is related to reduced working memory performance in older frontal and basal ganglia lesion patient populations^21,22^. In this study, we observed no difference in the amplitude or load-dependent modulation of CDA between younger and older adults. A previous study has reported alterations in CDA modulation in older adults^23^, but differences between this study and our present study are likely due to our study only presenting stimuli in one visual hemifield at a time. Thus, any age-related differences in the suppression of distractor processing were not tested, likely altering patterns of CDA modulation in older adults.

After memory array presentation, alpha amplitude in younger adults diverged between hemispheres, with ipsilateral amplitude higher than contralateral amplitude. Consistent with previous studies^3,4^, this alpha lateralization is suggestive of differential processing of the two visual hemifields and the deployment of selective spatial attention in anticipation of the test array, which participants knew would appear in the same visual hemifield as the memory array. This interpretation is also consistent with the lack of alpha lateralization in response to the spatially uninformative alerting cue. Compared to younger adults, older adults showed reduced alpha lateralization, as previously reported in studies with spatial cuing^12,24^. However, neither the degree of alpha lateralization nor the magnitude of CDA predicted older adults’ lower accuracy during load-three trials.

Instead, cue-evoked alpha phase resetting was less consistent in older adults and was predictive of behavioral performance even after adjusting for array-evoked alpha lateralization and delay-period CDA. Because the alerting cue appeared prior to any stimulus to be encoded in working memory, this result supports findings of reduced alertness in older adults^10,11^, with participants’ general attentional state being the single best predictor of accuracy more than a second later in the trial. While the age-related inconsistency in cue-evoked alpha phase resetting is opposite that in a previous study^24^, this discrepancy could be due to the lack of distractor stimuli and the briefness with which we presented the alerting cue (50 ms). This briefness potentially exacerbated any age-related alterations in cue response, which has not been observed in other studies^5,13^.

Interestingly, we also found that array-evoked ITC was similar between younger and older adults, despite previous reports showing increased ITC among older adults^24^. However, the large, asymmetric cue-evoked ITC differences between younger and older adults may have shifted the ITC baseline, artificially driving up younger-adult ITC. That is, the peak-to-peak difference between cue‐ and array-evoked ITC is much larger among older, compared to younger, adults. Nevertheless, that cue-evoked alpha phase consistency was predictive of behavioral performance is consistent with previous studies examining alpha phase resetting in response to task-relevant stimuli^25–27^. Our results extend these findings by demonstrating that alpha phase resetting in response to noninformative cues, even prior to presentation of to-be-remembered stimuli, can predict subsequent working memory performance.

Alpha phase prior to memory array presentation also predicted response time in high-load trials. This result provides further evidence for the effects of alpha phase on visual working memory^28^. These effects have also been demonstrated in visual detection paradigms^29,30^. Due to the consistent time interval between cue and memory array presentation in our study, it is possible that cue-evoked alpha phase resets led to subsequent memory array presentation at phases facilitative of or detrimental to perception or encoding of the memory array. Older adults’ inconsistency in phase response could have led to a greater number of instances in which memory array presentation occurred at suboptimal alpha phases, potentially explaining part of the age-related reductions in performance we observed during high-load trials. However, older adults’ weaker relationship between alpha phase and response time suggests age-related reductions in the influence of alpha phase on visual cognition. Physiologically, this reduced influence, as well as older adults’ inconsistent cue-evoked phase responses, may relate to age-related increases in neural noise^31,32^.

Overall, we find that oscillatory alpha dynamics may underlie age-related alterations in attention. Our analysis of alpha phase highlights reductions in older adults’ response and attentiveness to alerting cues, with such responsiveness being the strongest predictor of working memory performance. In addition, prestimulus alpha phase predicted performance on a trial-by-trial basis, but less reliably so in older adults. Given that lower performance in older adults can be explained by altered response to alerting cues prior to the task, age-related working memory decline is likely multifaceted and includes alterations in anticipatory attentional allocation as well as in stimulus encoding and maintenance. These findings suggest that changes in neural response, especially in older adults, can occur at multiple timepoints both before and after presentation of task-relevant stimuli, and such alterations likely all have an impact on later cognitive performance.

## Methods

### Behavioral Task

Healthy right-handed younger (20-30 year olds, *n* = 17, eight female) and older (60-70 year olds, *n* = 14, seven female) adults with normal or corrected-to-normal vision participated in a visual working memory paradigm. All participants gave informed consent approved by the UC Berkeley Committee on Human Research. In each trial, participants were instructed to maintain central fixation, and at the beginning of each trial, the central fixation cross flashed from gray to pink for 50 ms, alerting participants to the start of the upcoming trial (Fig. 1A). This alerting cue offered no information on either the size or location of upcoming visual stimuli. Three hundred ms after the end of the alerting cue, participants were presented with one, two, or three colored squares for 180 ms in only one visual hemifield. After a 900 ms delay period, during which time no stimuli other than the fixation cross were present, a test array of the same number of squares in the same spatial locations appeared. Participants would manually respond with their right thumb to indicate whether or not the test array had the same color squares as the initial memory array.

Behavioral accuracy was assessed using *d’*, a sensitivity measure that takes false alarm and miss rates into account to correct for response bias. To avoid mathematical constraints in the calculation of *d’*, we applied a standard correction procedure in the case of 100% hit rate or 0% false alarm rate. Specifically, hit rate was decreased to 1 - 1/(2*N*) when necessary, with *N* being the total number of trials. Similarly, false alarm rate was increased to 1/(2*N*) when necessary^33^.

### Data Acquisition

We recorded 64-channel scalp electroencephalography (EEG) from each participant. Participants were tested in a sound-attenuated EEG recording room using a 64+8 channel BioSemi ActiveTwo amplifier (Amsterdam, Netherlands). EEG was amplified (−3 dB at ~819 Hz low-pass, DC coupled), digitized (512 Hz), and stored for offline analysis. Horizontal eye movements (HEOG) were recorded with electrodes at both external canthi. Vertical eye movements (VEOG) were monitored with a left inferior eye electrode and either a superior eye or a fronto-polar electrode. All data was referenced offline to the average potential of two mastoid electrodes and analyzed in MATLAB^®^ (R2015A, Natick, MA) using custom scripts and the EEGLAB toolbox^34^.

### Data Preprocessing

EEG data was downsampled to 256 Hz and had DC offset removed. EEG data was then highpass filtered above 0.1 Hz using a two-way, fourth-order Butterworth infinite impulse response filter. Any channel whose standard deviation was ±2.5 standard deviations away from the mean standard deviation of all channels was spherically interpolated (on average, 2 channels per participant). Independent component analysis (ICA) was performed using the EEGLAB toolbox, and to remove blink artifacts, ICA components most correlated with the difference between the frontopolar and left inferior eye electrodes were removed.

For event-related potential (ERP) analyses and to detect trials with artifacts, continuous EEG data was lowpass filtered below 30 Hz using a two-way, fourth-order Butterworth infinite impulse response filter. Data was epoched around the onset of the memory array using a pre-stimulus baseline of −500 ms to −400 ms. For scalp topographic visualization, and to normalize electrode locations, electrode potentials were swapped right to left across the midline as though stimuli were always presented in the right visual hemifield, making left and right hemisphere channels contralateral and ipsilateral to the stimulus, respectively. Lateralized potentials were analyzed in this ipsilateral-contralateral fashion. Trials where the standard deviation of a scalp electrode exceeded three times the standard deviation of that electrode across all trials were excluded. For saccade trials, trials where the standard deviation of the difference between the HEOG channels exceeded three times the mean of the HEOG channels across all trials were excluded. On average, 69.6% of total trials or 165 trials were kept per participant. No participants were excluded.

### Data Analysis

Peak alpha frequency (PAF), the frequency of maximum power between 7 and 14 Hz, varies in a trait-like manner^35^ and predicts visual performance^36^. To estimate PAF for each participant, we constructed power spectral densities (PSDs) using Welch’s method. In order to account for individual differences in 1/*f* electrophysiological background, which changes with age^31^, we used robust linear regression to estimate and remove the slope and offset of log-log space PSDs prior to identification of PAF.

Continuous, non-lowpass-filtered EEG data was bandpass filtered with a 4-Hz passband centered on each participant’s PAF. Filters were designed as two-way finite impulse response filters with filter length equal to three cycles of the low cutoff frequency. For each channel, bandpass-filtered time series were converted to z-scores using the mean and standard deviation of artifact-free alpha-band data across all trials and conditions. After normalization, the absolute value and angle of the Hilbert transform of the continuous EEG data was used to extract alpha analytic amplitudes and instantaneous phases, respectively. The phase time series yields cosine phase values of (-π, π] radians, with π radians corresponding to the trough and zero radians to the peak of the oscillation. This method yields results equivalent to sliding-window fast Fourier transform and wavelet approaches^37^.

After epoching and removal of marked artifact trials, alpha analytic amplitude time series were subjected to event-related analyses, including the subtraction of baseline activity from −500 ms to −400 ms. To assess trial-to-trial phase consistency (also called intertrial coherence, ITC), event-related phase time series were extracted, and for each time point, the mean vector length of the timepoint’s phase distribution was calculated across trials (*circ_r.m* function in the CircStats toolbox^38^). This mean vector length represents the degree of ITC, with ITC of unity reflecting a single adopted phase across trials and a value of zero reflecting uniformly distributed phases across trials. To assess single-subject ITC significance, a resampling approach was used. For each participant, we randomly sampled *n*/2 trials and calculated time-resolved ITC. For each of these sub-samples, we then calculated mean cue/array-related minus mean baseline ITC value. This was done 1000 times to build a single-subject distribution of cue/array-related ITC strength, and *H_0_* is that the mean of the distribution of differences is zero. A one-sample, one-tailed *t*-test was used to compare the distribution of these differences against *H_0_* for each participant.

### Statistical Analyses

All analyses were performed on data from EEG channels O1/2, PO3/4, and PO7/8, with channels O1, PO3, and PO7 considered contralateral to the memory array. Multiple-factor statistical analyses were assessed via ANOVAs, with age as a between-group factor and memory load and hemisphere as within-group factors. Where sphericity assumptions were violated, degrees of freedom (and hence *p*-values) were adjusted using Greenhouse-Geisser corrections. All single-factor comparisons were analyzed via paired-samples or between-samples *t*-tests. For all alpha ITC analyses except those pertaining to single-subject ITC significance, ITC values were log_10_-transformed and baseline subtracted. Peak cue‐ and array-related ITC were assessed using the maximum ITC peak after cue and memory array presentation, respectively. To correlate circular variables like alpha phase with linear variables like response time, a circular-linear correlation was used (*circ_corrcl.m* function in the CircStats toolbox).

## Acknowledgements

We thank S.R. Cole, T. Donoghue, R. van der Meij, T. Noto, E.J. Peterson, and B. Postle for invaluable discussion and comments. This work was supported by the University of California, San Diego, Qualcomm Institute, California Institute for Telecommunications and Information Technology, Strategic Research Opportunities Program, and a Sloan Research Fellowship.

## Author Contributions

B.V. designed research. S.C.L., L.T., and B.V. performed research. T.T.T. and N.C.H. analyzed the data. T.T.T. and B.V. drafted the manuscript. All authors edited and approved the manuscript.

## Materials and Correspondence

Correspondence and requests for materials can be addressed to Tam T. Tran.

